# The neural basis for response latency in a sensory-motor behavior

**DOI:** 10.1101/623256

**Authors:** Joonyeol Lee, Timothy R. Darlington, Stephen G. Lisberger

## Abstract

We seek a neural circuit explanation for sensory-motor reaction times. We have found evidence that two of three possible mechanisms could contribute to reaction times in smooth pursuit eye movements. In the smooth eye movement region of the frontal eye fields (FEF_SEM_), an area that causally affects the initiation of smooth pursuit eye movement, neural and behavioral latencies have significant trial-by-trial correlations that can account for 40% to 100% of the variation in behavioral latency. The amplitude of preparatory activity, which represents the motor system’s expectations for target motion, shows negative trial-by-trial correlations with behavioral latency and could contribute to the neural computation of reaction time. In contrast, the traditional “ramp-to-threshold” model is contradicted by the responses of many, but not all FEF_SEM_ neurons. As evidence of neural processing that determines reaction time, the local field potential in FEF_SEM_ includes a brief wave in the 5-15 Hz frequency range that precedes pursuit initiation and whose phase is correlated with the latency of pursuit in individual trials. We suggest that the latency of the incoming visual motion signals combines with the state of preparatory activity to determine the latency of the transient response that drives eye movement.

## Introduction

When a major league baseball hitter faces a 95-mile-per-hour fastball, he has just over 400 ms from when the pitcher releases the ball to the time of contact with his bat. Extensive neural processing must occur in this short time to (1) process the visual inputs from the baseball, (2) decide whether or not to swing the bat, and (3) initiate a movement that is accurate in both location and timing. Timing is particularly important because a difference of a few milliseconds in the timing of the swing will determine whether or not the bat contacts the ball and whether the ball is driven into fair or foul territory. The temporal precision and accuracy of highly-skilled movements focuses attention on the neural mechanisms that decide exactly when to initiate a movement.

In the present paper, we ask what features of neural responses control behavioral latency using visually-guided smooth pursuit eye movements as a model system. Pursuit is driven by visual motion with latencies just under 100 ms, and is initiated in the rich form we study only in the presence of visual motion. The visual inputs that drive pursuit arise from extrastriate area MT (Newsome et al., 1988), where latencies to visual motion average around 60 ms (Lisberger and Movshon, 1999). Thus, only 40 milliseconds are available to transform visual motion signals into movement. We tend to think of the latency of pursuit in terms of the times required for visual signals to arrive in MT, traverse the cortex to the smooth eye movement region of the frontal eye field (FEF_SEM_), reach neurons in the pontine nuclei and the nucleus reticularis tegmenti pontis, and drive the simple-spike activity of Purkinje cells in the floccular complex of the cerebellum.

From first principles, however, multiple features of neural responses might control behavioral latency. One popular model posits that neural activity at a key site ramps up to a threshold that triggers movement. Faster ramps will reach threshold more quickly and trigger a movement with a shorter latency. This model appears to hold for saccadic eye movements (Hanes and Schall, 1996), and may apply for decision making, where one theory posits that ramping activity during the acquisition of sensory data accumulates evidence and leads to a movement to report a decision when the ramp crosses a threshold (Gold and Shadlen, 2007). However, ramp-to threshold may not apply for all kinds of movements. For example, as we found in area MT, the latency of the visually-driven transient response might be the key feature of neural responses that determines behavioral latency.

FEF_SEM_ is known from stimulation, lesion, recording, and anatomical studies to be causally involved in the generation of pursuit eye movements (Brodal, 1980; Gottlieb et al., 1994, 1993; Keating, 1991; Lisberger, 2010; MacAvoy et al., 1991; Ono and Mustari, 2009; Shi et al., 1998; Tanaka and Fukushima, 1998; Tanaka and Lisberger, 2001, 2002b, 2002a). It is a combined sensory and motor area and plays a role in setting the strength of the pursuit response based on previous experience and the signal-to-noise ratio of the sensory signal. The responses of neurons in FEF_SEM_ are rich enough to allow tests of multiple neural correlates of pursuit latency. The responses include both transient pulses of firing related to the initiation of pursuit and preparatory activity that evolves before a moving target initiates pursuit. Thus, we can test the ramp-to-threshold theory as well as the potential roles of the amplitude of preparatory activity versus the latency of the pursuit-related response in determining behavioral latency.

We find evidence for multiple computations that may control pursuit latency, with at best weak evidence in favor of the ramp-to-threshold theory. The trial-by-trial variation in the latency of pursuit-related FEF_SEM_ responses is linked closely to variation in the behavioral latency. Neural latency can account for 40 to 100% of the variation in behavioral latency. The amplitude of preparatory activity is negatively correlated with pursuit latency in many neurons, and could contribute to the latency computation. We also find a low frequency component in the local-field potential (LFP) that, itself, shifts in time (or phase) in relation to pursuit latency. We suspect that both the latency of the incoming visual motion signals and the expectation represented by preparatory activity contribution to neural and behavioral latency, and that the low-frequency LFP holds a key to understanding how they combine.

## Results

Our goal is to understand the neural circuit mechanisms that determine sensory-motor latency. Our analysis focuses on the trial-by-trial variation between neural and behavioral responses because trial-by-trial correlations provide a powerful tool for understanding the processing that occurs within a sensory-motor system. For example, our previous paper (Lee et al., 2016) implies that variation in the latency of responses in area MT could account for up to 70% of the trial-by-trial variation in the latency of pursuit.

Our strategy was to record local field potentials (LFP) and the spiking activity of multiple single neurons in the smooth eye movement region of the frontal eye fields (FEF_SEM_) while monkeys pursued a spot target that underwent step-ramp motion (Figure 1A, after Rashbass, 1961). Because we were interested in evaluating single-trial data and revealing trial-by-trial correlations between neural and behavioral responses, we typically collected more than 100 repetitions of a few target motions (Figure 1B).

**Figure 1.**
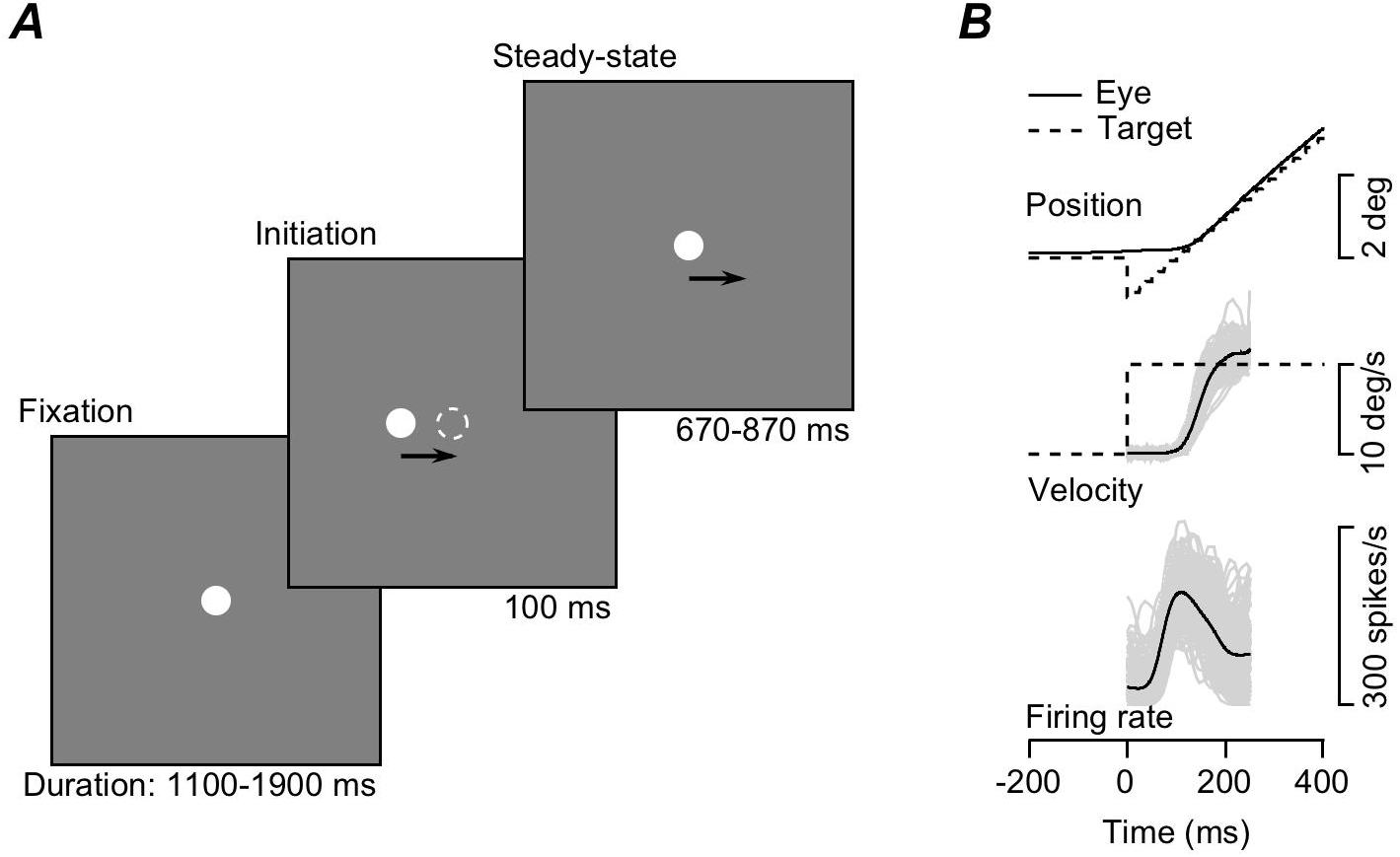
Structure of the task and example traces of step-ramp target motion and smooth pursuit eye movements. **A:** The three components of step-ramp target motion. **B:** Superimposed eye and target motion and spike density function traces. Dashed and continuous traces in the top two panels show target and eye motion. The black and gray traces for eye velocity and firing rate show averages across trials and data from individual trials.

Here, we present new results on the features of responses in FEF_SEM_ that contribute to neural and behavioral latency, and we compare our results in FEF_SEM_ with additional analyses of the data from prior recordings in area MT (Lee et al., 2016). We report three main findings. (1) The local field potentials in both FEF_SEM_ and MT include low-frequency components that are linked tightly to the latency of smooth pursuit eye movements and that are a signature of trial-by-trial correlations between the latencies of neighboring neurons. (2) The trial-by-trial variation in the latency of spiking responses in FEF_SEM_ is tightly correlated with the latency of pursuit eye movements, with the expected commensurate neuron-neuron latency correlations for pairs of FEF_SEM_ neurons (Schoppik et al., 2008). (3) The amplitude of preparatory activity is correlated negatively with behavioral latency, and could play a causal role in determining the latencies of FEF_SEM_ responses and pursuit eye movement.

### Trial-by-trial correlation between neural latency and pursuit latency

In response to target motion in their preferred direction, FEF_SEM_ neurons fire a transient of spikes followed by a sustained plateau, frequently at a lower rate (Figure 1B). As illustrated in Figure 1B, the onset of FEF_SEM_ responses usually precedes the onset of pursuit eye velocity, and both the firing rate response and the eye velocity of pursuit vary considerably in both latency and amplitude from trial-to-trial. Here, we leverage those variations to discover features of the neural computations that determine sensory-motor latency in the pursuit system.

Trial-by-trial correlations between the latency variations of FEF_SEM_ neurons and pursuit eye movements provide evidence that FEF_SEM_ controls sensory-motor reaction time to some degree. We used a procedure described elsewhere to estimate trial-by-trial latencies for pursuit eye movements (Lee et al., 2016; Lee and Lisberger, 2013) and to sort behavioral trials into 5 quintiles based on the estimated latency of pursuit in each trial (Figure 2A). We then revealed coordinated shifts in the latencies of eye movements and neural responses in FEF_SEM_ by averaging the time-varying firing rates in each quintile (Figure 2B).

**Figure 2.**
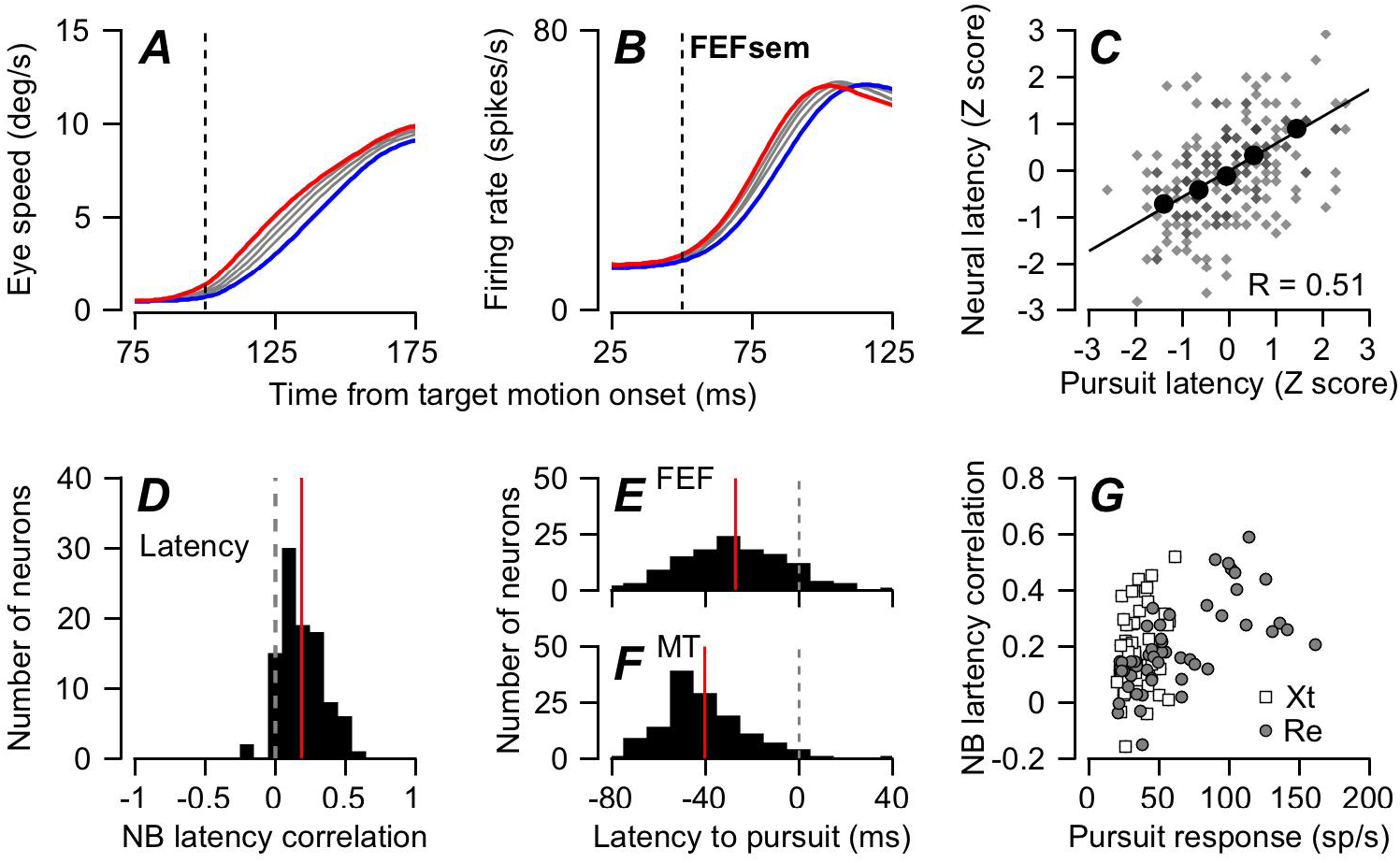
Trial-by-trial relationships between latency variation in pursuit behavior and firing of FEF_SEM_ neurons. **A:** Eye speed versus time during the initiation of pursuit. **B:** Firing rate of an example FEF_SEM_ neuron in the same intervals. In **A-B**, the five traces in each panel show data for quintiles of trials sorted by the latency of pursuit. Red and blue traces show data for the shortest and longest latency group. Gray colors show data for the middle groups. **C**: Plot of z-scored neural latency versus z-scored pursuit latency for an example neuron. Black circles show data for the 5 quintiles based on pursuit latency and small gray symbols show measurements for individual trials. Line was obtained by regression. **D:** Distribution of response latency-pursuit latency correlations for 99 FEF_SEM_ neurons. **E, F**: Distributions of latency from neural responses in MT and FEF_SEM_ to the onset of pursuit. **G**: Relationship between FEF_SEM_ neural response amplitudes and neuron-behavior latency correlations. The correlation between neural responses and neuron-behavior latency correlations was 0.46 (Pearson’s correlation, p = 1.4×10^−6^). Different symbols in **G** show data recorded in different monkeys.

To quantify the trial-by-trial latency correlations between FEF_SEM_ responses and behavior for each neuron, we applied the estimation method detailed in our previous publication (Lee et al., 2016). Briefly, we used an iterative Bayesian method to estimate the latency and gain of firing rate and eye velocity on each individual trial relative to a template created by averaging across groups of trials. We made these estimates on the basis of the trajectory of firing rate from −20 to 100 ms from the onset of neural response and of eye velocity from −20 to 80 ms after the initiation of smooth pursuit eye movement. By basing our estimates on the full transition of each variable from baseline to peak value, we ensure that our estimates of latency are based on the left-right shift of the full waveform relative to the average, and the estimate of response amplitude is based on the full transition from baseline to peak. We then obtained the Pearson’s correlation coefficient between the latencies of firing rate and eye movement by taking advantage of the fact that the correlation coefficient is defined by the regression slope of Z-transformed data (Figure 2C).

The correlation between neural and behavioral latency (Figure 2D) formed a distribution across all our neurons with a mean of 0.19 that was significantly different from zero (t-test p = 3.4×10^−22^, n=99 neurons). The mean correlation coefficients were 0.196 and 0.183 in the two monkeys individually and were significantly different from zero (t-test p = 7.13×10^−12^ and 1.46×10^−11^, n=50 and 49 neurons). In contrast, for the same sample of neurons, the correlation between the amplitude of the pursuit-related responses and behavioral latency had means of 0.007 and −0.010 in the two monkeys and were not significantly different from zero (t-test p = 0.61 and 0.61).

The absolute response latencies of FEF_SEM_ neurons (n=125) fit well within the expectations for a sensory-motor circuit. They preceded the initiation of pursuit by an average of 27 ms (Figure 2E) and followed the responses of MT neurons (n = 135) by an average of 13.3 ms (Figure 2F). The latency of responses in area MT neurons was 40.3 ms and the distributions of response latency for MT and FEF_SEM_ were statistically different (t-test, p = 4.2×10^−10^).

To verify that the small magnitude of neuron-behavior latency correlations was not an artifact of the vagaries of estimating single-trial neural latency from a handful of spikes, we evaluated the relationship between neural response amplitude and neuron-behavior latency correlation. Our premise was that artifacts of latency estimation would mitigate against large correlations in neurons with weak responses. However, we found that neurons with weak responses during pursuit could have either strong or weak neuron-behavior latency correlations while neurons with stronger responses had mainly strong neuron-behavior latency correlations (Figure 2G). Thus, we believe the values of neuron-behavior latency correlation.

### Local field potential in the 5-15 Hz range is linked to pursuit latency

Local field potential (LFP) provides an index of the neural processing within the localized region around the tip of the recording electrode. We reasoned that analysis of the LFP in our recordings should reveal correlates to confirm the phenomena we measured in spikes, and that the LFP might reveal special features of the neural computation that determines sensory-motor reaction time.

In FEF_SEM_, we find a fairly large trial-by-trial correlation between the phase of the LFP and the pursuit latency (Figure 3A). As outlined in the ***Methods***, we applied the Hilbert transform to LFPs that had been filtered with narrow pass bands at different frequencies and obtained instantaneous phase and power for each frequency and every millisecond in every trial. Then, we calculated the trial-by-trial correlation between pursuit latency and the instantaneous phase and power as a function of time. For FEF_SEM_, the correlation peaks for frequencies in the 5-15 Hz range and times around 100 ms after the onset of target motion, near the time of the initiation of pursuit. Analysis of LFP data in area MT (Figure 3B) revealed a similar correlation, albeit with a slightly smaller value of peak correlation in MT for the examples in Figures 3A and B.

**Figure 3.**
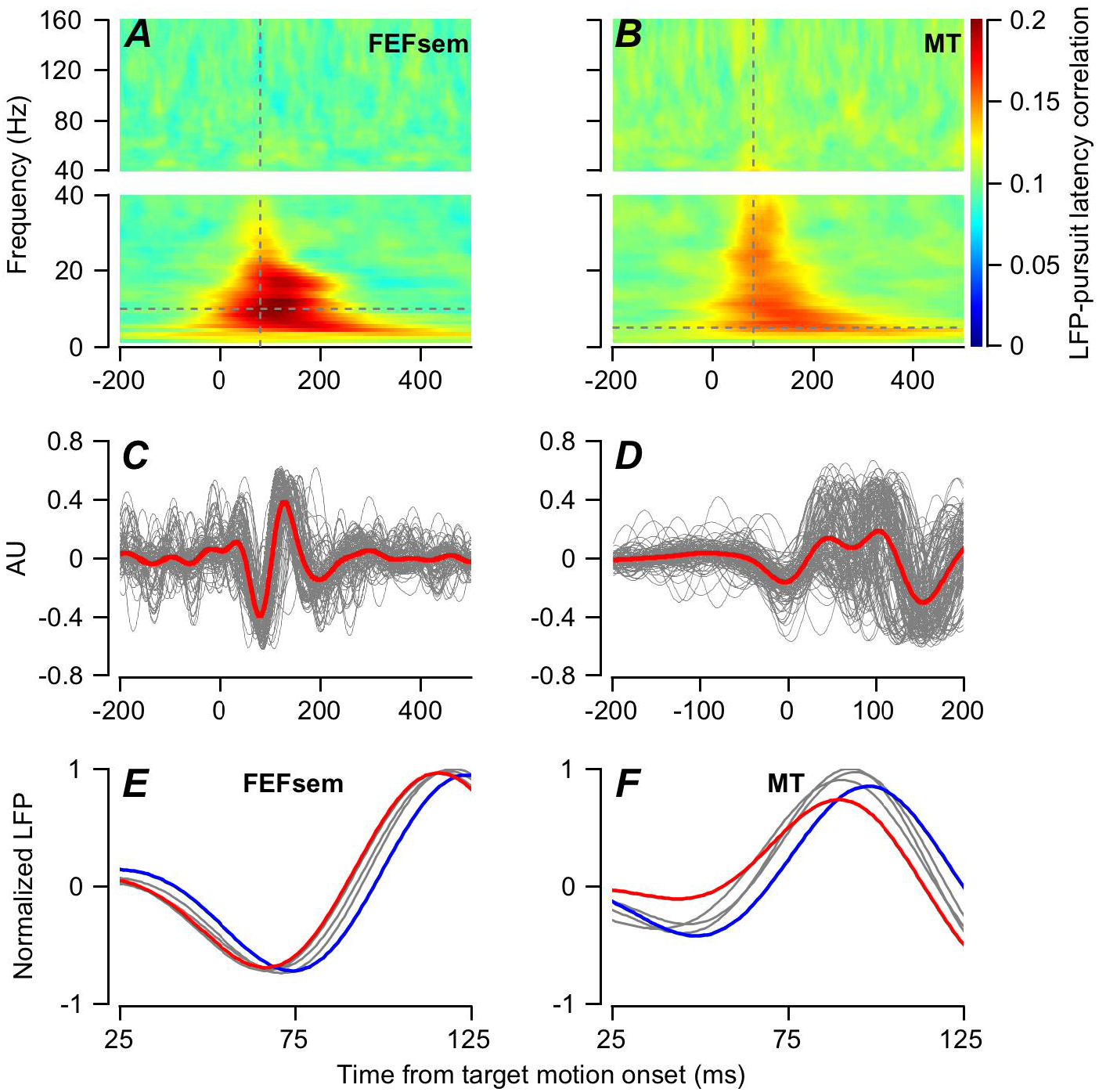
Local field potential and its correlation with the latency of pursuit. **A, B**: The color of each pixel plots the trial-by-trial correlations between pursuit latency and the phase of the LFP averaged across 123 sites in FEF_SEM_ (**A**) and 138 sites in MT (**B**) as function of LFP frequency in 1 Hz bands and time from the onset of target motion. **C, D**: Traces for local field potential filtered with a bandpass of 5-15 Hz for FEF_SEM_ (**C**) and MT (**D**). Red and gray traces show averages across all the recordings and data from individual average recordings for the FEF_SEM_. **E, F**: LFP sorted into quintiles according to pursuit latency, after normalizing individual filtered LFPs from individual trials by the maximum peak to trough distance. Red and blue traces show data for the shortest and longest latency group. Gray colors show data for the middle groups. Example neurons recorded in FEF_SEM_ (**E**) and MT (**F**).

The finding of a correlation between LFP phase and pursuit latency stimulated us to identify the component of the LFP that is linked to pursuit initiation. We band-pass filtered the raw LFP from each behavioral trial to extract the 5-15 Hz frequency component. When synchronized on the onset of target motion, the LFP in FEF_SEM_ from single trials contained a brief wave that started about 30 ms after the onset of target motion and underwent a negative-positive-negative fluctuation that ended about 200 ms after the onset of target motion. The shape of the LFP wave was consistent across the 123 LFP recordings as is shown in Figure 3C. In MT, the same analysis yielded a somewhat smaller and more complicated set of deflections, but otherwise the results in the two areas were remarkably similar (138 LFP recordings, Figure 3D). We next asked whether the LFP shifted in time in relation to pursuit latency by grouping the filtered LFP according to the quintiles of pursuit latency for each neuron. For two example neurons, this analysis revealed a relationship between the phase shift of the LFP and the latency of pursuit, both in FEF_SEM_ (Figure 3E) and MT (Figure 3F), consistent with the trial-by-trial correlations shown in Figures 3A and B.

Quantitative analysis of all recording sites from FEF_SEM_ and MT revealed that behavioral latency is much more strongly correlated with the phase shift of the LFP than with its power. Now, we correlated the behavioral latency on each trial with the power and phase in the LFP after filtering with a 5-15 Hz frequency window and plotted the results as a function of the time during the LFP. For an example LFP recorded in FEF_SEM_ (Figure 4A), the correlation between latency and LFP phase reaches a maximum correlation of ~0.6, while the correlation between latency and LFP power is smaller and negative. Population averages across 123 LFPs in FEF_SEM_ show the same effects (Figure 4C). In the population average, the correlation reaches a peak of 0.25 just more than 100 ms after the onset of target motion, which corresponds to the time of initiation of smooth pursuit. The results are similar for the LFP in area MT, although the effects are smaller (Figure 4B, D). We conclude that LFP in the 5-15 Hz frequency range is tightly linked to the latency of pursuit and may be related to neural processing in both MT and FEF_SEM_ that is contributing to sensory-motor reaction time. The differences in the details of the LFP wave in MT and FEF_SEM_ and its systematic relationship with behavioral latency strongly imply that the LFPs we found in the two areas are not simply an artifact of the procedures we used for filtering the LFP.

**Figure 4.**
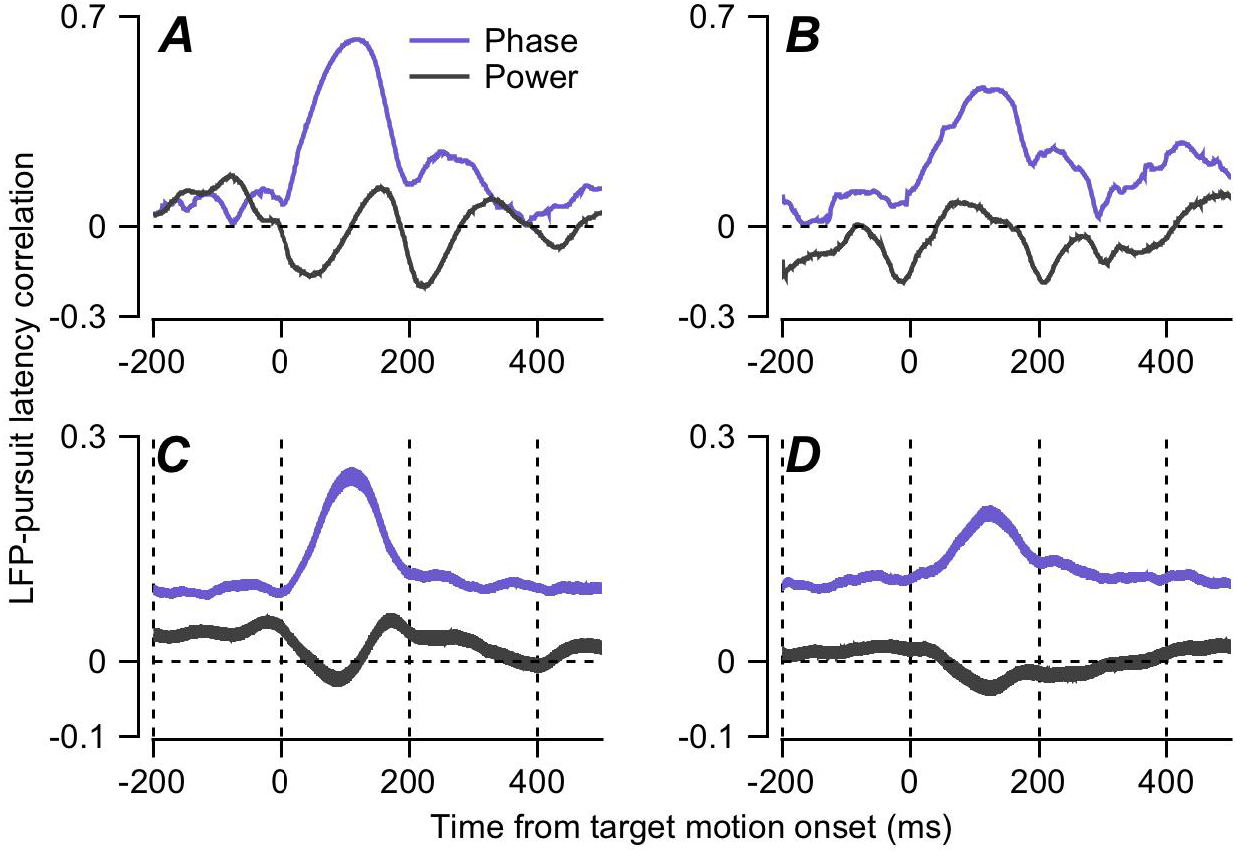
Quantitative analysis of the relationship between local field potential and pursuit latency. Graphs show example recordings in FEF_SEM_ (**A**) and MT (**B**) and averaged across all recording sites in FEF_SEM_ (**C**) and MT (**D**) as a function of the time during the wave. Blue and black traces show correlations of the LFP phase versus power, and the ribbons around the traces in **C** and **D** show the standard error of the mean across recordings.

### Relationships among LFP, neuron-behavior, and neuron-neuron latency correlations

The existence of neuron-behavior latency correlations in FEF_SEM_ implies that each neuron acts as a proxy for many other neurons with correlated response latencies. The predictive power of the latency of a single neuron results from the causal effect of it and its partners. The relationship between LFP and response latency supports this interpretation. To better understand the neural processing that creates these relationships, we next explored the interactions among neuron-behavior latency correlations, neuron-neuron latency correlations, and LFP.

The magnitude of each neuron’s neuron-behavior latency correlation is correlated with the coherence of its spikes with the LFP. This observation supports the conclusion that responses in any individual neuron are related to responses in multiple neighbor neurons. For the two example neurons in Figures 5A and B, one (A) has a neural latency that is weakly correlated with the behavioral latency, while the other (B) has neural latency that is strongly correlated with the behavioral latency. The spike-field coherences of the two neurons differed the most in 5-15 Hz frequency range when we then plotted the z-scored spike field coherences as a function of frequency at 100 ms after the onset of target motion (Figure 4C).

**Figure 5.**
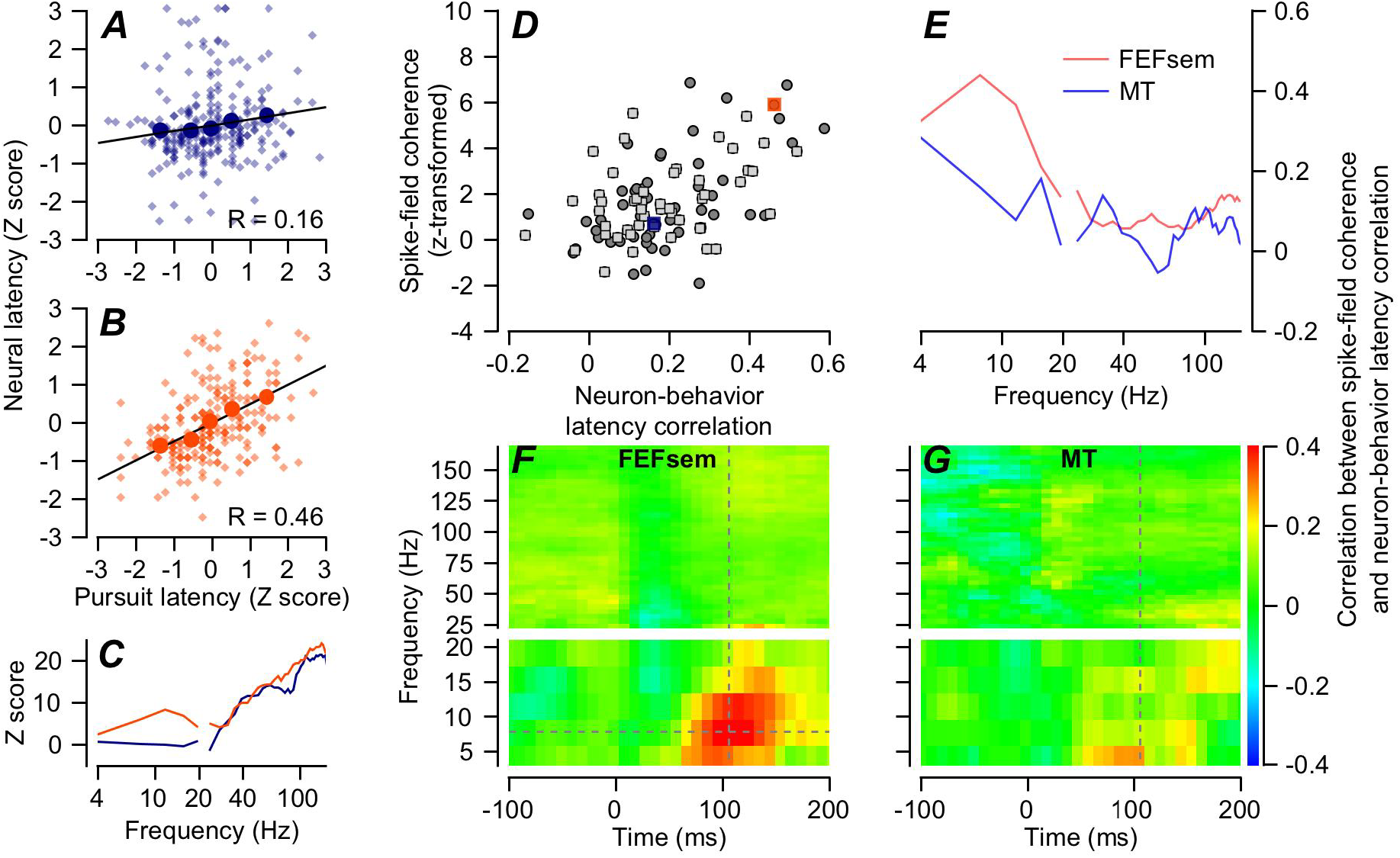
Relationship between spike-field coherence and FEF_SEM_-pursuit latency correlation. **A-B**: Plots of z-scored neural latency versus z-scored pursuit latency for 2 example neurons. Navy blue and orange circles show data for the 5 quintiles based on pursuit latency and smaller diamond symbols show measurements for individual trials. Lines were obtained by regression. FEF_SEM_-pursuit latency correlation of the neurons in panels **A** and **B** are low versus high. C: Z-scored spike-field coherences of the two example neurons as a function of frequency measured at a time window centered at 106 ms from motion onset. Colors match **A** and **B**. **D**: Each symbol shows data for a different neuron and the scatter plot shows the correlation between z-scored spike-field coherences and FEF_SEM_-pursuit latency correlations. Orange and navy-blue squares are data of example neurons. Dark gray circles and light gray squares are data from monkeys Re and Xt. **E**: Red and blue lines show the correlation between z-scored spike-field coherences and neuron-behavior latency correlations measured at a time window centered at 106 ms averaged across all recording sites in FEF_SEM_ and MT. **F, G**: The color of each pixel shows the correlation between z-scored spike-field coherence and neuron-behavior latency correlations for example recording sites in FEF_SEM_ (**F**) and MT (**G**), plotted as a function of frequency in the coherences and time from the onset of target motion. The intersection of the dashed lines in **F** shows the time and frequency used for panel **D**.

To quantify this relationship for all FEF_SEM_ neurons, we plotted the z-scored spike-field coherence in a time window centered at 106 ms and a frequency window centered at 7.8 Hz versus the neuron-behavior latency correlation, revealing a strong relationship (Figure 5D). The Pearson’s partial correlation (controlling for firing rate) of 0.43 and 0.36 for monkeys Re and Xt (p = 0.0023 and 0.012, n=50 and 49 neurons) suggests that the LFP in the frequency range from 5-15 Hz represents an aspect of neural processing in FEF_SEM_ that is related in some way to behavioral latency.

Reanalysis of the neural data collected from area MT for our earlier paper (Lee et al., 2016) reveals that the relationship between the magnitude of spike-field coherence and neuron-behavior latency correlation is weaker in MT than in FEF_SEM_, and the peak in the frequency range from 5-15 Hz is broader in MT (Figures 5E). The colormap in Figure 5F confirms that the forgoing analysis used the correct frequency band and time interval. Here, each pixel shows the correlation coefficient from graphs like that in Figure 5D for all times and frequencies. The relationship between neuron-behavior latency correlation and LFP is strongest in the frequency range of 5-15 Hz in the time interval from 80-150 ms after the onset of target motion. Comparison with Figure 5G confirms that the relationship between neuron-behavior latency correlation and the LFP is much weaker in MT. These differences support our suggestion that the significance of the 5-15 Hz LFP in FEF_SEM_ is a product of the underlying biology and not simply an artifact of the signal processing, which was identical in the two areas.

The relationships among neuron-behavior latency correlations, behavioral latency, and the low frequency LFP component suggest that the neural synchronization with the LFP could be linked to synchrony in the responses of FEF_SEM_ neurons, that is to their degree of neuron-neuron latency correlation. We verified this prediction by measuring neuron-neuron latency correlations between pairs of FEF_SEM_ neurons and comparing them with the neuron-behavior latency correlations (Figure 6A, B). We selected 93 pairs of simultaneously recorded neurons that satisfied criteria outlined in the ***Methods***, and calculated trial-by-trial latency correlation following the methods we published before (Lee et al., 2016). Pairwise neuron-neuron latency correlations had a distribution with a mean of 0.04 (Figure 6A). The Pearson correlation is 0.53 between neuron-neuron latency correlation and the product of the neuron-behavior latency correlations for the two neurons in the pair (Figure 6B, p=7.25×10^−19^, p=1.2×10^−4^ and 1.5×10^−4^ for each monkey individually), consistent with expectations if neuron-neuron latency correlations cause neuron-behavior latency correlations. We also found a strong correlation (Pearson’s ρ=0.4, p=2.23×10^−4^) between neuron-neuron latency correlation and the mean of the spike-field coherences for the pair of neurons under study (data not shown).

**Figure 6.**
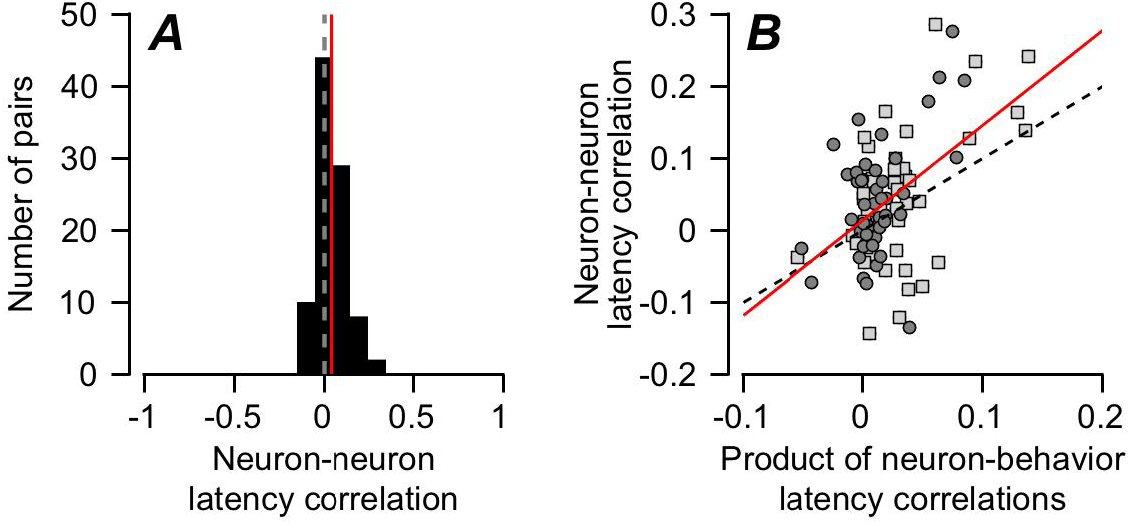
Neuron-neuron latency correlations in pairs of FEF_SEM_ neurons. **A:** Distribution of neuron-neuron latency correlations between pairs of FEF_SEM_ neurons. Vertical red line shows the mean correlation. **B:** Each point plots data for a different pair of FEF_SEM_ neurons, and the scatter plot shows the relationship between the product of FEF_SEM_-pursuit latency correlations and neuron-neuron latency correlations. The dashed line has a slope of one and the red line is a regression fit. Dark gray circles and light gray squares are data from monkeys Re and Xt.

### Other features of neural responses that may contribute to pursuit latency

It makes sense that the correlated latency of the transient responses of neurons in the FEF_SEM_ plays a role in determining the latency of pursuit initiation. But, what mechanisms control the neural latencies in FEF_SEM?_ Several possibilities exist: (1) the correlated latency of visual motion inputs from the population response in MT (Lee et al., 2016); (2) the time when a ramp increase in firing rate reaches a threshold (“ramp-to-threshold”, Hanes and Schall, 1996); and (3) the amplitude of preparatory activity that ramps up during fixation before the onset of target motion (Darlington et al., 2018). We used analyses of trial-by-trial correlations with behavioral latency to evaluate the variables that might affect neural latencies in FEF_SEM_.

We observed trial-by-trial correlations between the duration of fixation and behavioral latency and between the amplitude of preparatory activity and behavioral latency, supporting the possibility that the amplitude of preparatory firing rate at the time of target motion onset could contribute to neural latency in FEF_SEM_. The time from the onset of the fixation point to the start of target motion varied between 1100 and 1900 ms in these experiments, and the trial-by-trial correlation between the duration of fixation and the latency of pursuit was negative (n=131 neurons, average Pearson’s correlation across 153 sessions, r=-0.18, t-test p=2.68×10^−47^), meaning that longer fixation is associated with shorter behavioral latency. The correlations in the data in the present paper using a 2-or 4-direction pursuit task agreed well with those from the same monkeys in an earlier paper that used a single-direction pursuit task (Figure 7A, C) (ρ=-0.238, t-test p=2.25×10^−130^, n=95 sessions with 4 speed/contrast conditions, Darlington et al., 2018).

**Figure 7:**
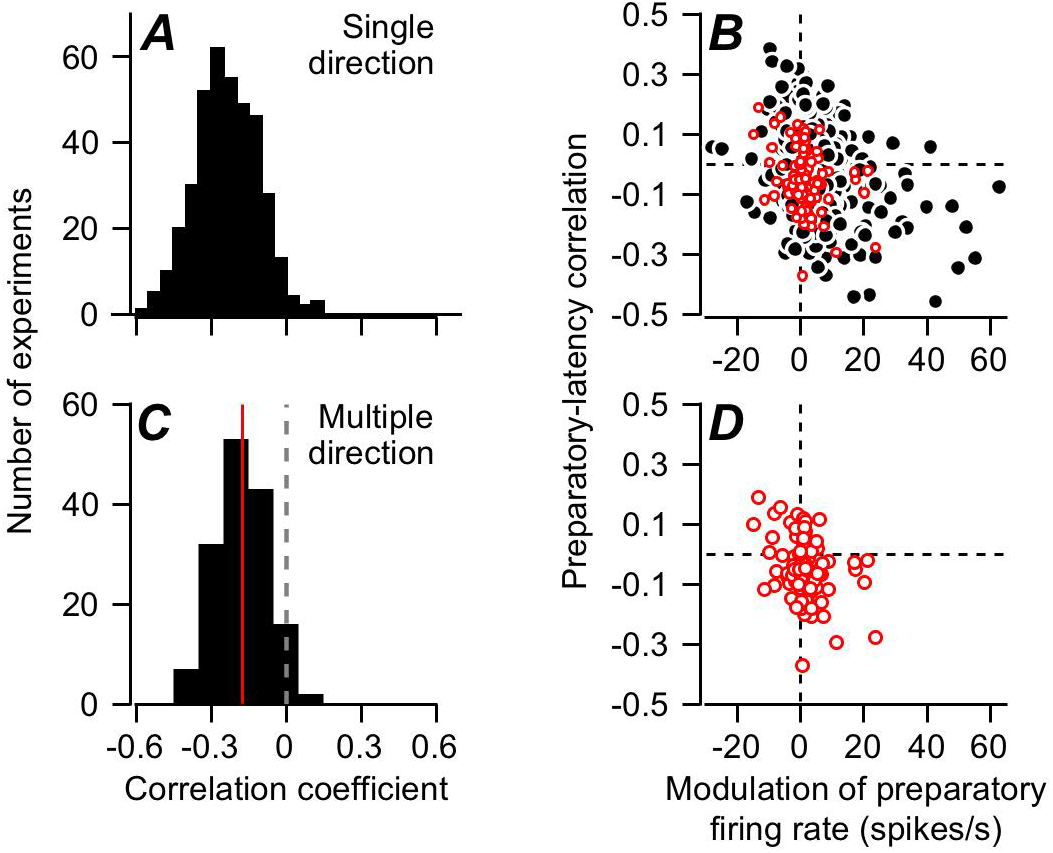
Relationship between preparatory activity and behavioral latency. **A, C:** Distributions of trial-by-trial correlations between duration of fixation and pursuit latency. **A** reanalyzes previously-reported single-direction experiments in monkeys Xt and Re, while **C** reports results for the experiments in this paper, from the same 2 monkeys but for blocks of trials with 2 or 4 directions of randomly-interleaved target directions. **B, D:** Scatter plots showing the trial-by-trial correlation of preparatory activity at the end of fixation with pursuit latency versus the modulation of preparatory activity across fixation. Each symbol shows results for a different experiment. Again, **B** shows data for previous single-direction experiments and **D** shows data for multiple-direction experiments in the present paper. Red symbols in **D** have been replotted in **B**.

The correlation between behavioral latency and firing rate at the end of fixation averaged −0.04, but was quite large for neurons with large modulation of preparatory activity during fixation before the onset of motion. The correlation between the amplitude of preparatory modulation and the preparatory-latency correlation (Figure 7D) was −0.32 (Pearson’s correlation, p=0.001, n=99 neurons), and the neurons from our multiple-direction blocks of trials agreed well with those from single-direction experiments (ρ=-0.32, t-test p=4.55×10^−9^, n=321 directions collected across 164 neurons, Figure 7B). The wider range of modulation of preparatory activity in the single-direction experiments could reflect either the monkey’s greater certainty about the impending target motion, or a difference in the population of sampled neurons.

We conducted two analyses as explicit tests that failed to support the ramp-to-threshold theory of movement latency for FEF_SEM_.

> In the first analysis, we divided all the trials for a single condition of target motion in a single neuron into quintiles according to the latency of pursuit in each trial and computed the time-varying average of firing rate within each quintile. We analyzed the firing rate in the interval from 20-40 ms before the behavioral latency in each quintile (Figure 8A), plotted the firing rate as a function of the latency, and fitted the data with a regression line (Figure 8B). The ramp-to-threshold theory posits that the slope of the regression line for these plots should be zero. Across two full populations of FEF_SEM_ neurons, the slopes were close to zero for neurons with relatively weak pursuit-related responses and were strongly positive for neurons with stronger pursuit-related responses (Figure 8C). These two populations of neurons were recorded from the same two monkeys using a single-direction pursuit task with a high-contrast random dot patch target moving at 20 deg/s (black symbols) or a 2-or 4-direction task (red symbols) with a spot target moving at 10 deg/s.
>
> In the second analysis, we computed the time-varying trajectories of mean eye velocity and firing rate separately for target motions at 2, 10, and 20 deg/s at high and low contrast for each individual neuron. We then measured the average behavioral latency and the firing rate in the interval from 20-40 ms before the onset of pursuit, plotted the firing rate as a function of the latency. Again, the ramp-to-threshold theory predicts slopes of zero, and we found mostly positive slopes for neurons with large pursuit-related responses and again slopes near zero for neurons with smaller responses. Together, these results do not speak strongly for the ramp-to-threshold theory for the FEF_SEM_, especially since the neurons with slopes of zero showed little activity in the interval from 20-40 ms before the onset of pursuit (Figure 8B).

**Figure 8.**
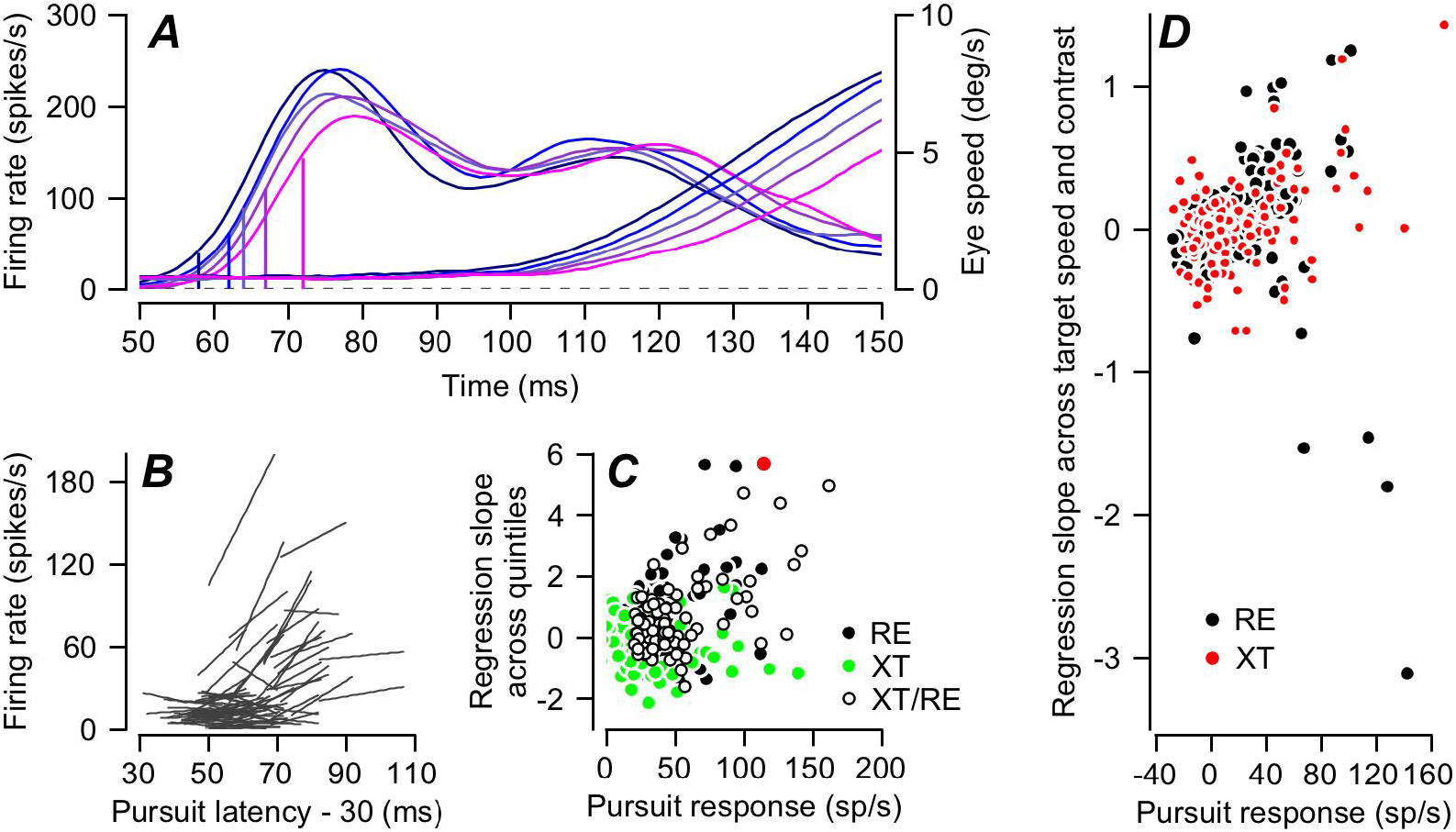
Direct tests of the ramp-to-threshold hypothesis for individual neurons in FEF_SEM_. **A:** Firing rate (left-hand traces) and eye velocity (right-hand traces) for an example neuron divided into quintiles according to pursuit latency in individual trials and plotted as a function of time from the onset of target motion. The 5 vertical lines on the firing rate traces show the values of firing rate 30 ms before the latency of eye velocity in each quintile. **B**: Linear regression lines fitted to each FEF_SEM_ neuron’s plots of firing rate 30 ms before the onset of pursuit versus pursuit latency for the 5 quintiles of pursuit latency. Each line shows data for a different neuron. **C**: Slope of the regression line versus amplitude of the pursuit-related response. A slope of zero would support the ramp-to-threshold hypothesis. Each symbol shows a different neuron. Open symbols show data from this paper, red symbol shows the neuron in **A**, and filled green and black symbols show results of the same analysis for our earlier single-direction experiments on the same two monkeys. **D**: Same plot as **C**, but now using data solely from our earlier single-direction experiments and plotting the regression slopes for points taken from targets of high-or low-contrast moving at 2, 10, or 20 deg/s.

### Contribution of the correlated latency variation in FEF_SEM_ on trial-to-trial variation of smooth pursuit latency

Our data document relationships among a low-frequency component of LFP, neuron-behavior latency correlations, and neuron-neuron latency correlations of FEF_SEM_ neurons. These results indicate that the latency variation of the FEF_SEM_ neural population, which arises from the correlated latency variation between neurons, has an impact on behavioral latency variation. Therefore, we asked how much of smooth pursuit latency variation can be accounted for by the correlated neural latency variation in FEF_SEM_ neurons.

We followed the same procedure that we used previously for estimating the contribution of the correlated latency variation of area MT neurons to latency variation in the initiation of smooth pursuit (Lee et al., 2016). The main feature of the procedure is that we need to estimate what we call the “underlying probability of firing” and compute realistic correlations between the latencies of the underlying probabilities of spiking for pairs of model FEF_SEM_ neurons so that we can generate properly correlated model spike trains (details below). We might think of the underlying probability of firing in terms of the summed post-synaptic current in each FEF_SEM_ neuron.

Our analysis has multiple steps.

1. We calculate the underlying probability of firing from the spiking data for each FEF_SEM_ neuron by averaging spike density across multiple repetitions of the same target motion.
2. We estimate the standard deviation (SD) of the latency of the underlying firing probability by assuming different values of SD, assembling 100 trials with the underlying firing probability shifted in time according to the SD, laying down simulated spikes with realistic discharge irregularity for each specific neuron, and performing our analysis of the latency of spike trains to estimate the SD of spiking latency. Then, we find the SD of the underlying spiking probability that matches the SD of spiking probability we measured from the actual data through the linear regression function between SD of underlying firing probability and estimated SD of simulated spikes. Figure 9A shows the relationship between SD of spiking and SD of underlying firing probability from our sample of FEF_SEM_ neurons.
3. We use a similar procedure to estimate the correlation between the neural latency variation of the underlying firing probability of spiking and the behavioral latency variation. Again, we create model neurons with different degrees of correlation between the latencies of the underlying probability of spiking and of pursuit, lay down simulated spikes with realistic discharge irregularity for each neuron in our sample, and performed our analysis of the trial-by-trial correlation between the latencies of the simulated spike trains and the behavior. We then use the neuron-behavior latency correlation for each neuron’s actual spikes to estimate its neuron-behavior latency correlation for the underlying probability of firing (Figure 9B). The average neuron-behavior latency correlations were 0.19 for spikes and 0.3 for the underlying probability of firing.
4. We estimate neuron-neuron latency correlations between underlying probabilities of spiking of pairs of simultaneously recorded neurons using the same procedure outlined above, except with the underlying probability of firing for pairs of neurons. The range of actual correlations is wider for the underlying probability of firing versus the spikes (Figure 9D versus C), and neither shows a compelling relationship to the latency difference between the two neurons. Neuron-neuron latency correlations for spiking and the underlying probability of spiking averaged 0.057 and 0.19.
5. Using model parameters based on the distribution of parameters derived for our sample of FEF_SEM_ neurons, we simulated a population of 1000 spiking FEF_SEM_ neurons. The model neurons had realistic time dependent shapes of underlying firing probability of spiking, coefficient of variation (CV) of inter-spike interval, neural latency SD, and neuron-neuron latency correlations. We ran the simulation 200 times to represent the responses to 200 trials of target motion. For each trial, we created a population spike density function by convolving each spike train with a Gaussian function having an SD of 10 ms and then averaging across the responses of 1000 model neurons. We decoded behavioral latency from the population spike density function using the same procedure we had used for estimating latency from the smooth pursuit data.

**Figure 9.**
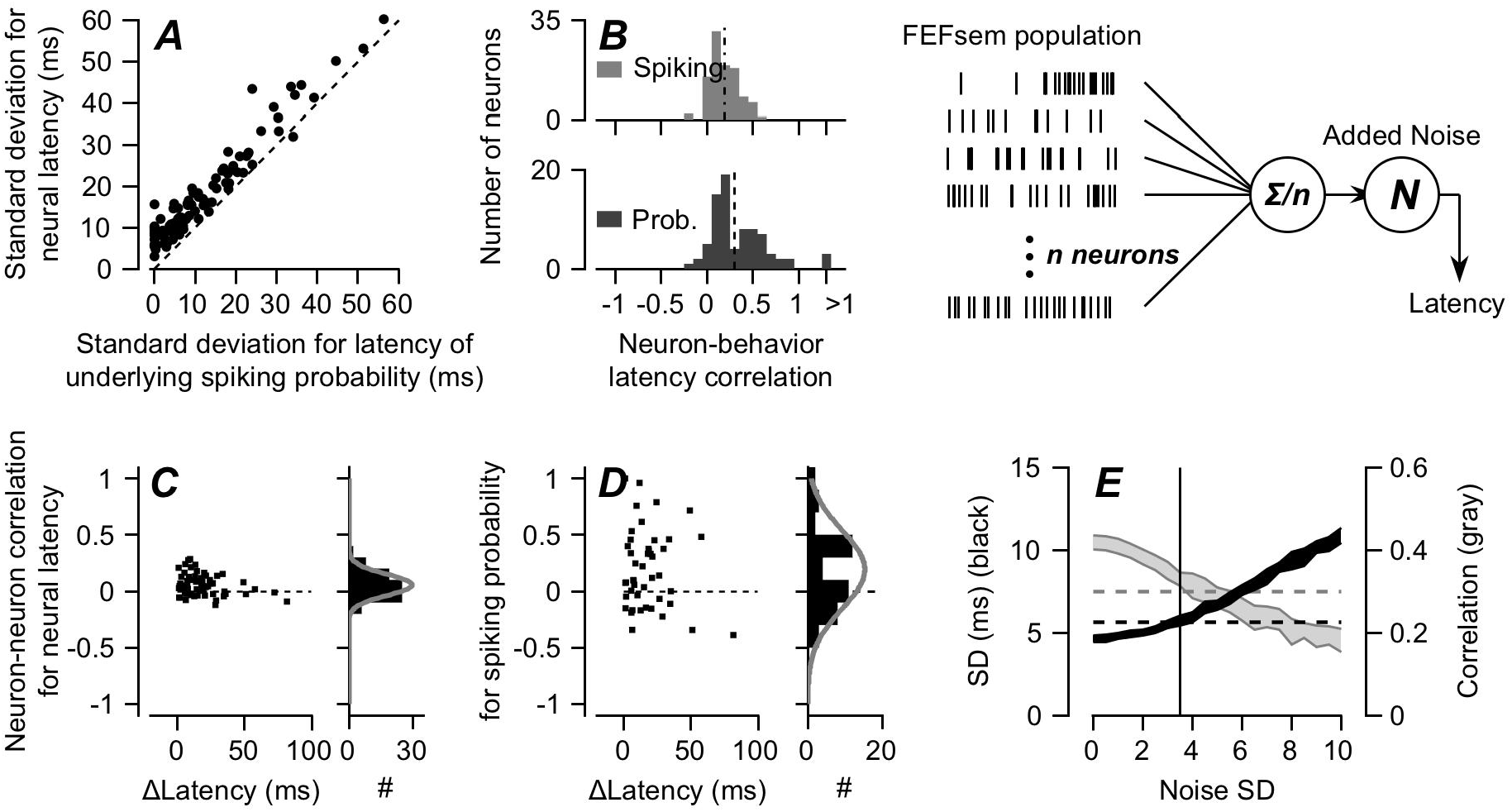
Computational analysis for the effect of neuron-neuron latency correlation on behavioral latency variation. **A**: Relationship between latency SD of firing rate and latency SD of underlying firing probability of spiking. Each symbol shows one neuron. Dashed line is the unity line. **B**: Distributions of FEF_SEM_-pursuit latency correlations for firing rate and for underlying probability of spiking. The number of neurons included on this analysis is different from the previous figures. **C**: Distribution of neuron-neuron latency correlation for firing rate as a function of the mean latency differences between neurons. **D**: Distribution of neuron-neuron latency correlation for underlying probability of spiking in relation to mean neural latency differences. **E**: Predictions for pursuit behavior based on decoding a model FEF_SEM_ population for different values of downstream noise. Black and gray filled areas show standard deviation of latency and neuron-behavior latency correlation averaged across 10 simulations. Vertical line shows SD of 3.5 ms for Gaussian noise added downstream.

For the model FEF_SEM_ population, the predicted standard deviation of behavioral latency was 4.6 ms in the absence of downstream noise and increased as a linear function of Gaussian noise applied downstream (Figure 9E). Therefore, variation in the latency of responses in FEF_SEM_ can account alone for at least 40% of the nominal behavioral latency variation SD of 10 ms (Osborne et al., 2005). In the experiments reported here, the average SD of pursuit latency was 5.6 ms (Figure 9E, black dashed line), so that FEF_SEM_ could actually be contributing almost all of the behavioral latency variation.

In the model FEF_SEM_ population, the predicted neuron-behavior latency correlation for underlying firing probability averaged 0.42 in the absence of downstream noise and decreased as a function of the noise added downstream increased (Figure 9E, gray dashed line). In our data, the neuron-behavior latency correlation for the underlying probability of firing in our FEF_SEM_ neurons averaged 0.3. When we added random noise from a Gaussian distribution with SD of 3.5 ms (Figure 9E, vertical line), the SD of predicted behavioral latency was 5.7±0.3 ms and the predicted neuron-behavior latency correlation for underlying firing probability was 0.33±0.02, both comparable to what we observed in the data. We conclude that under certain assumptions, the latency variation in FEF_SEM_ could account for all of the behavioral variation in pursuit latency.

## Discussion

We think that FEF_SEM_ has a causal role in determining pursuit latency because of its known causal relationship with pursuit eye movements (Brodal, 1980; Gottlieb et al., 1994, 1993; Keating, 1991; Lisberger, 2010; MacAvoy et al., 1991; Ono and Mustari, 2009; Shi et al., 1998; Tanaka and Fukushima, 1998; Tanaka and Lisberger, 2001, 2002b, 2002a). Our analysis reveals four important findings that help us to understand how FEF_SEM_ controls sensory-motor reaction time. (1) Latency variation of the pursuit-related response in FEF_SEM_ is correlated with latency variation of pursuit behavior and could account for 40-100% of the variation in behavioral latency. (2) The amplitude of preparatory activity could contribute to the regulation of neural and behavioral latency, but the data do not support the ramp-to-threshold theory strongly for pursuit eye movements. (3) When a neuron’s latency variation is highly correlated with the pursuit latency variation, the neuron’s latency variation is likely to be correlated with that of neighboring neurons. (4) The latency of pursuit and the correlated variation of neuronal latencies in the FEF_SEM_ both are linked tightly to the local field potential in the 5-15 frequency range, and this feature of the LFP seems likely to reflect essential features of the neural control of latency. We propose two components that contribute to response latencies in FEF_SEM_: the state of preparation and the timing of visual motion-related inputs. Preparatory and visual motion signals combine to determine the timing of the pursuit-related FEF_SEM_ responses that are linked to pursuit reaction times.

### Control of pursuit latency

The control of pursuit latency appears to use a neural mechanism quite different from that for the control of saccade latency. For pursuit, the timing of the overall envelope of firing rate in FEF_SEM_ (present data) and MT (Lee et al., 2016) encodes the latency of the impending pursuit eye movement. The ramp-to-threshold theory that seems to work for saccadic eye movements in other parts of the frontal eye fields (Hanes and Schall, 1996), but not for pursuit eye movements in FEF_SEM_. However, the trial-by-trial correlation between the amplitude of preparatory activity and the latency of pursuit raises the possibility that preparatory activity plays a role in determining the latency of the pursuit-related responses in FEF_SEM_ and therefore also of pursuit itself.

We have now reported significant trial-by-trial neuron-behavior latency correlations in three major nodes for smooth pursuit eye movement: extrastriate visual area MT, FEF_SEM_, and the floccular complex of the cerebellum. The presence of the 5-15 Hz LFP in both MT and FEF_SEM_ suggests that they aren’t completely independent, but simultaneous recordings from MT and FEF_SEM_ will be required to determine the extent to which FEF_SEM_ and MT control pursuit latency jointly or independently. We admit that our current understanding of the role of FEF_SEM_ in pursuit eye movements somewhat clouds the question of its contribution to pursuit latency. Our correlation analysis indicates that the outputs from MT and FEF_SEM_ both contribute to pursuit initiation, while prior data indicate that FEF_SEM_ modulates the strength of visual-motor transmission from MT, potentially without actually driving pursuit eye velocity itself. We do not know how a multiplicative modulation would affect pursuit latency. However, one possible answer is that the gain control signal emanating from FEF_SEM_ is critically important in determining pursuit latency because the visual motion signal from MT cannot initiate the movement when gain is zero.

### Origin of correlated variation of FEF_SEM_ neural latency and pursuit latency

Our data imply that the correlated latency variation across neurons in FEF_SEM_ contributes to the actual trial-by-trial variation in the latency of pursuit and leads to the impressive neuron-behavior latency correlations we recorded in FEF_SEM_. Thus, the same framework seems to apply for correlated latencies and response amplitudes in many areas of the brain, including area MT for smooth pursuit eye movements (Hohl et al., 2013; Huang and Lisberger, 2009; Lee et al., 2016; Lee and Lisberger, 2013), area MT for perceptual decisions (Shadlen et al., 1996), FEF_SEM_ for pursuit eye movements (Schoppik et al., 2008), and subcortical vestibular neurons (Liu et al., 2012) and area MSTd (Gu et al., 2014) for heading discrimination. The relationship between the size of the neuron-behavior correlations for a given neuron and its spike-field coherence with the LFP supports strongly the conclusion that neural activity acquires trial-by-trial predictive power for behavioral variation because the neuron under study is part of a population of neurons with correlated variation.

Noise correlations across the relevant population is a necessary, but not sufficient, condition to create neuron-behavior correlations in either latency or response amplitude. In addition, the amount of behavioral variation added downstream from the area under study must be small, because downstream noise will inevitably reduce neuron-behavior correlations (Schoppik et al., 2008). In the neural circuit for pursuit eye movements, it is striking that downstream noise does not eliminate neuron-behavior correlations in either latency or response amplitude. We conclude, as we did before (Osborne et al., 2005), that the trial-by-trial variation in pursuit latency and amplitude is a consequence of correlated noise in cortical areas, and that sub-cortical components of the circuit loyally follow the instructions given by the cortex without judging their veracity. Indeed, the presence of neuron-behavioral correlations limits the amount of noise that can be injected downstream, challenging to some degree the conclusions of Bakhtiari and Pack (2019) about how much noise is created by noise correlations in MT versus downstream from MT.

### The possible meaning of 5-15Hz LFP

We found relationships among neuron-neuron latency correlation, neuron-behavior latency correlation, and spike-field coherence in 5-15 Hz frequency range. The three measures covary across neurons that we recorded: a neuron with high synchrony with 5-15 Hz LFP has high neuron-neuron latency correlation and high behavior-neuron latency correlation. Their covariation implies that the three factors originate from one mechanism.

The 5-15 Hz frequency range of the LFP overlaps with the alpha range of neural oscillation, but we do not think this is a meaningful piece of information. We find it challenging to reconcile the linkage between the LFP wave and pursuit initiation with the relationship of alpha wave to feedback signaling (Michalareas et al., 2016), attention (Saalmann et al., 2012), attentional gating (Foxe and Snyder, 2011) or working memory (Gevins et al., 1997). We also note but cannot reconcile our findings with the LFP suppression in the beta range that precedes arm movement (Best et al., 2017), or the correlation between beta power and reaching reaction time in the primary motor cortex (Khanna and Carmena, 2017). A previous study reported a traveling beta wave in macaque area V4 that is correlated with the parameters of saccadic eye movements (Zanos et al., 2015), including an interesting phase relationship between beta frequency LFP and the timing of saccade offset. Our findings differ from this study in the specific frequency of LFP (beta vs. 5-15 Hz), the behavior (saccade vs. smooth pursuit), the brain area (area V4 vs. FEF_SEM_), and the time epochs for the correlation (after saccade offset vs. before pursuit initiation). In many ways, the predictive value of the LFP wave component in FEF_SEM_ for neural and pursuit latency resembles that for gamma synchrony in the LFP in area MT for neuron-neuron spike count correlations (Lee and Lisberger, 2013). It is curious that latency and spike rate are related to LFP in such different frequency ranges.

Finally, we are struck by the fact that the LFP wave in the 5-15 Hz frequency range shifts in time in parallel with pursuit latency, and that its phase is so well correlated with pursuit latency on a trial-to-trial basis. We view the 5-15 Hz frequency components in the LFP as a signature of critical neural activity that gets synchronized in the FEF_SEM_ to facilitate initiation of a pursuit eye movement. It could reflect internal activity in FEF_SEM_, could be a correlate of a trigger for pursuit initiation from another node in the pursuit circuit, or might be simply a consequence of latency correlations in inputs from area MT that report the sensory estimate of target speed and direction. We suspect that understanding the latency-related component of the LFP will prove to be a key in unraveling that neural mechanisms that determine behavioral latency in our particular sensory-motor system, and possibly in other systems as well.

## Materials and Methods

Two adult male rhesus monkeys were trained to pursue a circular dot that executed a step-ramp trajectory of target position (Rashbass, 1961), in exchange for juice or water reward. Before training, we performed two separate surgeries using sterile procedure. In the first surgery, a head holder was implanted on the skull for head restraint, and a stainless-steel chamber was implanted on the skull, over the cross-section of arcuate sulcus and central sulcus. In the second surgery, a scleral search coil was implanted in one eye (Ramachandran and Lisberger, 2005). After finishing behavioral training, we performed a craniotomy inside the chamber for introducing quartz-insulated platinum/iridium electrodes into the brain. All experiments were conducted at Duke University using methods that had been approved in advance by the *Institutional Animal Care and Use Committee* at Duke University. Methods conformed to the *National Institutes of Health Guide for the Care and Use of Laboratory Animals*. The latency distribution for neurons and the relationship between spikes and LFP in area MT were obtained by reanalyzing the data reported in previous papers (Lee et al., 2016; Lee and Lisberger, 2013). We also performed a few analyses on data collected from the same 2 monkeys in previous experiments on FEF_SEM_ (Darlington et al., 2018).

### Data acquisition

Visual stimuli were presented on a gamma-corrected 24-inch CRT color monitor (Sony Trinitron), whose spatial resolution was 2304 by 1440 pixels and vertical refresh rate was 80 Hz. The monitor was placed 60 cm from the monkey and the screen covered 44 by 29 deg of horizontal and vertical visual fields. Accurate measurement of visual motion onset timing is guaranteed by the post-hoc data collection using photodiode. Since we did not use the photodiode system during recording experiments, all the visual motion onset timing has been corrected by the reference data collected after the experiments. Horizontal and vertical eye position and velocity were sampled at 1 kHz. Eye velocity was obtained by differentiating and low-pass filtering eye position at a cut-off frequency of 25 Hz using an analog circuit.

During experiments, we slowly lowered one to three electrodes or tetrodes in the FEF_SEM_ and recorded spikes and local field potentials (LFP) using the Tetrode Mini-matrix System (Thomas Recording GmbH). The input impedance and gain of the Mini-matrix preamplifier were 1GΩ and ×19 individually. The input impedance of the preamplifier was sufficiently higher than the impedance of the single electrodes and tetrodes that we used (Thomas Recording GmbH, 2-4 MΩ and 1-2 MΩ individually), ensuring that our recordings would be relatively free from phase distortion in low frequency LFPs (Nelson et al., 2008). For LFPs, signals were low-pass filtered with a cut-off frequency of 170 Hz and digitized at a sampling rate of 2 kHz. For action potentials, signals were high-pass filtered with a cut-off frequency of 150 Hz and those that crossed a preset threshold were digitized at a sampling rate of 40 kHz. Digitization, filtering, and amplification of signals were done with a Plexon MAP system (Plexon Inc.).

We isolated single neural spikes online, using a window discriminator for analyzing and displaying neural response properties. We performed more accurate off-line discrimination of single neural responses later, using spike waveforms that exceeded a certain threshold. We took exceptional care in spike sorting because any significant sorting errors might work as an added noise in trial-by-trial correlation analysis. First, we performed initial spike sorting using principal component analysis. Then, we refined the sorting by visually inspecting the waveforms in a smaller time chunk, to detect any waveform changes caused by the slow drift of electrodes. We used Plexon Offline Sorter (Plexon Inc.) for all the offline spike sorting procedure. Sorted spikes were converted to time stamps with a time resolution of 1 ms and were inspected again visually to look for obvious sorting errors.

### Experimental design

We recorded neural activity in FEF_SEM_ while two rhesus monkeys were engaged in smooth pursuit eye movement tasks. We identified FEF_SEM_ using functional and anatomical markers. We implanted a recording chamber over the arcuate sulcus so that we could have perpendicular access to the rostral bank of the sulcus. As we advanced electrodes, we first looked for the area that shows oculomotor responses. We then advanced the electrodes past saccade-related neurons until we found neurons that were strongly modulated by smooth pursuit eye movements. Then we isolated single neuron responses and performed two experiments.

In the first experiment, we tested the direction tuning of the neurons under study during smooth pursuit eye movement. We randomly interleaved saccade trials and pursuit trials, where saccade trials required the monkeys to make 8 directions (45 deg interval) of 15 deg amplitude saccade eye movements and pursuit trials required them to make 8 directions (45 deg interval) of pursuit eye movements at 15 deg/s. We used a white circular dot as a saccade target and random dot stimulus and/or grating as pursuit targets. The random dot stimulus was made from white and black dots so as for us to control luminance contrast of the stimulus (see Yang et al., 2012). Typically, luminance contrast of the random dot stimulus was 100% and contrast of the grating stimulus was 12%. We usually used both stimuli, but either sufficed for determining the direction tuning of FEF_SEM_ neurons. All the visual stimuli were presented on a gamma corrected, gray background. We obtained saccade and pursuit direction tuning curves online using a least-square fitting method and used the tuning curves to choose the preferred pursuit direction of the neurons under recording, and to test whether the neurons were more-strongly tuned to smooth pursuit or saccadic eye movement. Only the 133 neurons with larger responses during pursuit than during saccades were admitted for further analysis.

In the second experiment, we recorded neural responses while monkeys tracked a yellow circular dot that moved in a step-ramp trajectory. Typically, we choose two pursuit directions that provided good compromises among the preferred directions of the neurons under study and one target speed (10 deg/s or 20 deg/s); sometimes we used both target speeds and more than two directions. The size of the step in the step-ramp target motion was 1 and 3 deg for target speeds of 10 and 20 deg/s. After the experiment, we screened all trials visually and excluded from further analysis any pursuit trials that contained saccades or microsaccades in the interval from 100 ms before to 210 ms after target motion onset.

### Estimation of pursuit latency and FEF_SEM_-pursuit latency correlation

We used an objective method that was developed in our previous study to quantify the latency of pursuit for each trial(Lee et al., 2016; Lee and Lisberger, 2013). Briefly, we computed the average horizontal and vertical eye velocities across trials and used the averages from 20 ms before to 80 ms after the initiation of pursuit as templates. Here, we used the initial 80 ms of smooth pursuit, instead of 100 ms to include more trials in the analysis. The overall result for pursuit latency was the same whether we used 80 ms or 100 ms for the analysis. We shifted and scaled the templates in every trial and estimated the best match using the least-square method. The best time shift defined the pursuit latency in each trial. We included a trial in the further analysis only if the best-fitted estimate accounted for more than 80% of eye-velocity variance. We proceeded to further analysis only if 80 or more trials passed this test.

We estimated the response latency of the average responses of FEF_SEM_ neurons through a statistical method. First, we obtained spike density functions by convolving spike trains with a Gaussian window of a 10 ms standard deviation and averaging the individual spike density functions across trials. Then, we set a baseline by averaging neural activity in the interval ±25 ms from motion onset. If the average PSTH between 30 ms and 300 ms from motion onset exceeded the baseline activity plus 20% of the standard deviation of the baseline and continued to do so for more than 50 ms, the first time bin of the 50 ms was selected as the latency of the average firing rate.

We estimated trial-by-trial FEF_SEM_-pursuit latency correlations using a method developed in our previous study (Lee et al., 2016). First, we excluded any neurons whose responses in the interval from 1 to 100 ms after neural response latency failed to reach 20 spikes/s, leaving 99 FEF_SEM_ neurons from the two monkeys for further analysis. Then, we selected one pursuit condition per neuron that provided the largest average response across the interval from 1 to 100 ms after the time chosen as the neural response latency. By taking advantage of the fact that the correlation coefficient is a regression slope of Z-transformed data (Rodgers and Nicewander, 1988), we obtained the correlation by taking a geometrical mean of the two regression coefficients – one coefficient with pursuit latency as independent variable and neural latency as dependent variable, and the other with neural latency as independent variable and pursuit latency as dependent variable.

To calculate a regression coefficient when pursuit latency is the independent variable, we divided the trials from each neuron into five equal-sized groups (quintiles) according to the estimated pursuit latency. We computed the spike density functions by averaging across the trials in each quintile of trials. Using the mean spike density function for all trials as a template, we estimated the best fitting values of time shift for each quintile by shifting and scaling the template to match the quintile’s spike density. Then, we calculated the regression coefficient for the estimated time shift of the mean spike density as a function of the mean pursuit latency for each quintile.

To calculate a regression coefficient when neuronal latency is an independent variable, we used an iterative process to estimate neuronal latency in individual trials. Using the mean spike density function for all trials as the template, we estimated time shifts and scaling factors for the spike density from each individual trial. We then updated the template using a Bayesian procedure (Bollimunta et al., 2007) and repeated the estimation procedure two more times. Then, we divided trials into quintiles according to the final estimates of neural latency and computed mean spike density function for each quintile of trials. We estimated the five best fitting values of time shifts by shifting and scaling the template along the mean spike density function of each quintile. Finally, we calculated the regression coefficient for pursuit latency as a function of the estimated neural latencies for the 5 quintiles. We obtained the Pearson’s correlation coefficient by taking geometrical mean of the two regression coefficients. We applied the same procedure using the responses of pairs of neurons to estimate neuron-neuron latency correlations. We started from 134 unique pairs, and again selected pairs with the average of the two average responses across the interval from 1 to 100 ms after each neural latency exceeds 20 spikes/s, which leaves 93 pairs in the further analysis.

### Local field potential analysis

#### Spike-field coherence

We preprocessed the LFP data with a Butterworth filter to remove 60 Hz line noise, using a function in the Fieldtrip MATLAB toolbox (Oostenveld et al., 2011). Then we analyzed the ‘spike-field coherence’ between the spikes and the LFP recorded from the same electrode. We used the Chronux MATLAB toolbox (Bokil et al., 2010) for all the spike-field coherence analysis. For the low frequencies (1-20Hz), we used a Hanning window as a taper. For frequency ranges between 20 and 170 Hz, we used five spheroidal tapers with 15 Hz of spectral smoothing. We calculated the spike-field coherence in 150 ms sliding windows with a step size of 10 ms. We calculated coherence and transformed the coherence using the variance stabilization method (Bokil et al., 2007). To correct a potential confound that can occur at the low frequencies due to the transient responses (Jarvis and Mitra, 2001), we shuffled trials and calculated the variance stabilized coherences 1000 times and generated the “null distributions” of coherences across frequencies. Then we obtained a z-score as a function of frequency using the shuffle analysis as in our previous study (Lee and Lisberger, 2013).

#### LFP phase-pursuit latency correlation

First, we used a 6^th^ order Butterworth filter to bandpass filtered the LFP and isolated frequency in bandwidths of 2 Hz and 1 Hz steps from 0 to 160 Hz. We obtained power and phase information from each bandpass filtered component by applying Hilbert transformation. Then, across trials and for each millisecond within trial, we calculated the correlation between instantaneous phase/power at that instant and estimated pursuit latency (Figures 2A, B). From the correlation map, we selected the frequency range that maximized the correlation, isolated the LFP in the 5-15 Hz range using a 6^th^ order Butterworth filter, and calculated the correlation between instantaneous phase/power and pursuit latency after Hilbert transformation. Because phase information has a circular form and pursuit latency has a linear form, we used the correlation coefficient calculation for circular and linear data (Berens, 2009).

### Underlying probability of spiking and computer simulations

As we showed in our previous study (Lee et al., 2016), variation in neural latency has two components, one created by variation in the timing of the underlying probability of spiking, and the other created by the stochastic nature of spike timing. To estimate the SD of the latency of the underlying probability of spiking, we used a computational simulation approach separately for each FEF_SEM_ neuron. (1) We calculated the coefficient of variation (CV) using inter-spike intervals in a time window from 100 to 300 ms after the average neural latency. (2) We recalculated the peristimulus time histogram to estimate a template for the underlying probability of spiking by aligning each trial according to its estimated neural latency and applying a Gaussian filter (3 ms SD) to the re-aligned PSTH. (3) In different runs where the SD of the latency variation was an independent variable, we jittered the template of underlying probability in latency across trials using random values from a Gaussian distribution with a mean of zero and SD chosen from predetermined values (1, 2, 4, 6, 8, 10, 12, 16, 20, 24, 28, 32, 36, 40). (4) For each run, we used the neuron’s CV as the basis for creating simulated spike trains that follow a renewal Gamma process (Brown et al., 2002; Koyama and Shinomoto, 2005). (5) We estimated latency SD from the simulated spike train using the same procedure that we used for real data. (6) We used the simulated relationship between latency SD for spikes and underlying probability of firing to estimate the actual latency SD of the underlying probability given the latency SD measured from each neuron’s spiking. In this step, we included neurons only if the regression explained more than 40% of the variance, reducing the number of neurons included in the further analysis from 99 to 94 (Figure 7A).

Once we estimate SD of latency for spiking and underlying firing probability, we can estimate latency correlation between underlying probability and behavior or between two underlying probabilities.

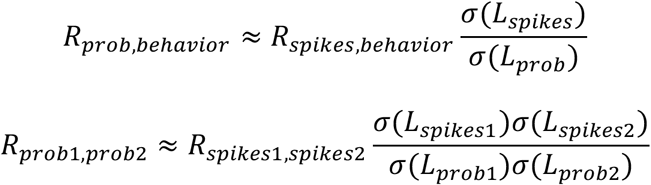

Where *L* represents latency. Following the two equations, we estimated latency correlation between underlying probability and behavior, and latency correlation between the two underlying probabilities. When we estimated latency correlation between underlying probability and behavior, we only selected neurons whose estimated latency SDs of underlying probability of spiking are the same or bigger than one, to avoid over-estimation of the correlation. This resulted in sample reduction for the estimation of latency correlation between underlying probability and behavior, from 94 to 76.

#### Computer simulations

To estimate how much of pursuit latency variation could originate from FEF_SEM_, we conducted a computational analysis. We simulated a population of 1000 spiking FEF_SEM_ neurons with responses and correlation structure that is statistically the same as our sample of FEF_SEM_ neurons. We used re-aligned time-varying mean spike density functions of our FEF_SEM_ recordings to determine the underlying probability of spiking. We also randomly selected the latency SD of the underlying firing probability of spiking from a Gamma distribution fitted to our observed distribution of latency SD with a shape parameter к of 1.39 and scale parameter θ of 10.16. For each model neuron, we selected a value of SD from the Gamma distribution, drew 200 latencies from a Gaussian distribution with that SD, and shifting the underlying firing probability of spiking in time by those values for our 200 simulated trials. In the population of 1000 model FEF_SEM_ neurons, we also retained the structure of the neuron-neuron latency correlations for the underlying probability of firing derived from our data: the distribution of pairwise latency correlation followed a Gaussian distribution with a mean of 0.19. For each trial in each model neuron, we modeled the inter-spike interval distribution by using the shape of the latency-jittered time dependent underlying firing probability to scale a renewal Gamma process based on the CV of inter-spike interval distribution in our data (Brown et al., 2002; Koyama and Shinomoto, 2005). We estimated the spike density of each simulated neuron by applying a Gaussian filter with SD of 10 ms and computed the population average across the 1000 simulated neurons’ spike density functions in each trial. From these population spike density functions, we estimated the latency in individual trials. We simulated the added downstream noise by generating random number that follows a Gaussian distribution, with zero mean and SD ranged from 0 to 10 ms in 0.5 ms step. We repeated the whole simulation process 10 times to calculate the mean and SD of simulated neuron-behavior latency correlation for the underlying firing probability and SD of simulated latencies.

## Acknowledgments

We thank S. Tokiyama, S. Ruffner, and S. Happel for technical assistance. We also thank Dr. J. Patrick Mayo for help in visual stimulus timing calibration. Research supported by NIH grant R01-EY027373 (SGL) and F30-EY027684 (TRD). Joonyeol Lee was partly supported by IBS-R015-D1.

The authors declare no competing financial interests.

## Author contributions

Conceptualization, J.L and S.G.L.; Methodology, J.L. and S.G.L.; Investigation, J.L. and TRD; Formal Analysis, J.L. and TRD; Writing – Original Draft, J.L. and S.G.L.; Writing – Review and Editing, J.L., T.R.D, and S.G.L.; Funding Acquisition, S.G.L.

